# Integral-Omics: serial extraction and profiling of metabolome, lipidome, genome, transcriptome, whole proteome and phosphoproteome using biopsy tissue

**DOI:** 10.1101/2024.09.27.614689

**Authors:** Wei Li, Jing Sun, Rui Sun, Yujuan Wei, Junke Zheng, Yi Zhu, Tiannan Guo

## Abstract

The integrative multi-omics characterization of minute amount of clinical tissue specimens has become increasingly important. Here, we present an approach called Integral-Omics, which enables sequential extraction of metabolites, lipids, genomic DNA, total RNA, proteins, and phosphopeptides from a single biopsy-level tissue specimen. We benchmarked this method with various samples and applied the workflow to perform multi-omics profiling of tissues from six patients with colorectal cancer and found that tumor tissues exhibited suppressed ferroptosis pathway at multi-omics levels. Together, this study presents a methodology that enables sequential extraction and profiling of metabolomics, lipidomics, genomics, transcriptomics, proteomics and phosphoproteomics using biopsy tissue specimens.

## Introduction

Disease complexity arises from synergistic effects and intricate regulation of biomolecules ^1, 2^. Multi-omics is crucial to understand disease mechanisms at the molecular level ^3-5^. A key requirement for integrated multi-omics analysis is the extraction of various biomolecules from the same sample. Typically, disease-related multi-omics studies divide a sample into several parts to extract the needed components. ^6-9^. For tissue-based multi-omics, a common approach involves homogenizing a bulk tissue specimen (typically >100 mg) into a fine powder, which is then aliquoted for further analysis ^10, 11^. However, this strategy is limited to relatively large tissue specimens, such as those from late-stage tumors. For smaller biopsy-level specimens (<20 mg), especially from early-stage diseases, developing an integrated methodology for sequential extraction of multiple biomolecules from a single tissue specimen is crucial ^10, 12-14^.

A few studies have endeavored to perform multiple omics using the same tissue specimen. Lipidomics and metabolomics co-extraction methods have become highly robust, primarily relying on the biphasic extraction techniques developed by Folch *et al.* (methanol/chloroform/water) and Matyash *et al.* (methyl tert-butyl (MTBE)/methanol/water), with subsequent modifications by researchers ^14-19^. Some researchers have extended these co-extraction methods to include proteomics by solubilizing proteins from precipitates after metabolite and lipid extraction. For instance, Burnum-Johnson *et al.* ^20^ isolated lipids, metabolites and proteins from a single pathogen sample based on the Folch’s method, and the Coon lab ^18^ employed the Matyash’s method to isolate lipids, metabolites and proteins from 1700 worms.

Some researchers have also expanded the co-extraction of metabolism, lipids, protein and nucleic acid. Leuthold *et al.* used the *mir*Vana^TM^ miRNA Isolation Kit (Thermo Fisher Scientific) and acetonitrile to isolate RNA and metabolites from 4-50 mg kidney tissues, but the lysis buffer’s reducing agents, salts, and detergents caused matrix effects and ion suppression during metabolite detection ^21^. Woodward *et al.* sequentially isolated metabolites, lipids, and RNA from 10-40 mg of brain tissue using methanol/chloroform/water and the *mir*Vana™ miRNA Isolation Kit, effectively addressing the ion suppression issues noted by by Leuthold *et al* ^22^. Roume *et al.* isolated metabolites, RNA, DNA and protein from single sample using the Folch’s method with AllPrep DNA/RNA/Protein Mini Kit (Qiagen) ^23^. In 2020, Hasegawa *et al.* built on the work of Roume *et al.* to refine the metabolite extraction process ^24^.

They incorporated ethylenediaminetetraacetic acid (EDTA), butylated hydroxytoluene (BHT), and potassium chloride (KCl) into the Folch’s extraction method and utilized the RNeasy MinElute Cleanup Kit (Qiagen) for the purification of small RNAs. This methodological enhancement successfully isolated polar and non-polar oxylipin metabolites, DNA, RNA, small RNA, and protein from about 30 mg of porcine brain tissue. However, these methods all rely on commercial reagent kits. Additionally, researchers have extracted DNA, RNA, and proteins from 10 million cancer cells using thiocyanate guanidine and cesium chloride, a method that bypasses commercial kits but requires an ultracentrifuge, making it complex and time-consuming ^25-27^. Current sequential multi-omics methods are limited by the number of layers they can extract or the need for commercial kits, specialized instruments, and none have demonstrated robust performance with biopsy-level (≤10 mg) tissue specimens.

We present Integral-Omics, a novel multi-omics workflow for serial extraction of metabolites, lipids, genomic DNA, total RNA, proteins, and phosphopeptides from as little as 10 mg of fresh tissue. The method employs a modified biphasic extraction (MTBE/methanol/DEPC water), followed by gentle ice bath ultrasound for biomolecule release and parallel extraction. We benchmarked Integral-Omics against independent extractions and a commercial kit, then demonstrated its applicability to biopsy-level clinical samples, including colorectal cancer specimens. This workflow shows promise for multi-omics characterization of small biomass clinical biopsy samples.

## Results

### Overview and rationale of the Integral-Omics workflow

We reviewed and integrated well-established methods for extracting metabolites, lipids, genomic DNA, RNA, proteins, and phosphopeptides. After designing the extraction sequence, comprehensively testing reagent compatibility, and comparing the methods, we developed a cohesive workflow called ‘Integral-Omics’ (**Figure 1**). Mouse liver was utilized as the test sample due to its homogeneity ^28^. The workflow (**Figure 1**, **Supplementary Note 3**) firstly uses the modified Matyash method (MTBE/methanol/DEPC H_2_O) to extract lipidome and metabolome. Subsequently, through a gentle ice bath sonication method, RNA and proteins are further released into the modified 8 M urea lysis buffer (100 mM ammonium bicarbonate (ABB), DEPC H_2_O), while the DNA is extracted from the precipitate using phenol/chloroform/isoamyl alcohol. The supernatant is divided into two aliquots, with 150 μL used for RNA extraction by TRIzol™ and 100 μL subjected to tryptic digestion of proteins. These volumes can be adjusted based on the specific requirements of the study. After tryptic digestion, peptides are further cleaned and subjected to phosphopeptide enrichment.

**Figure 1.**
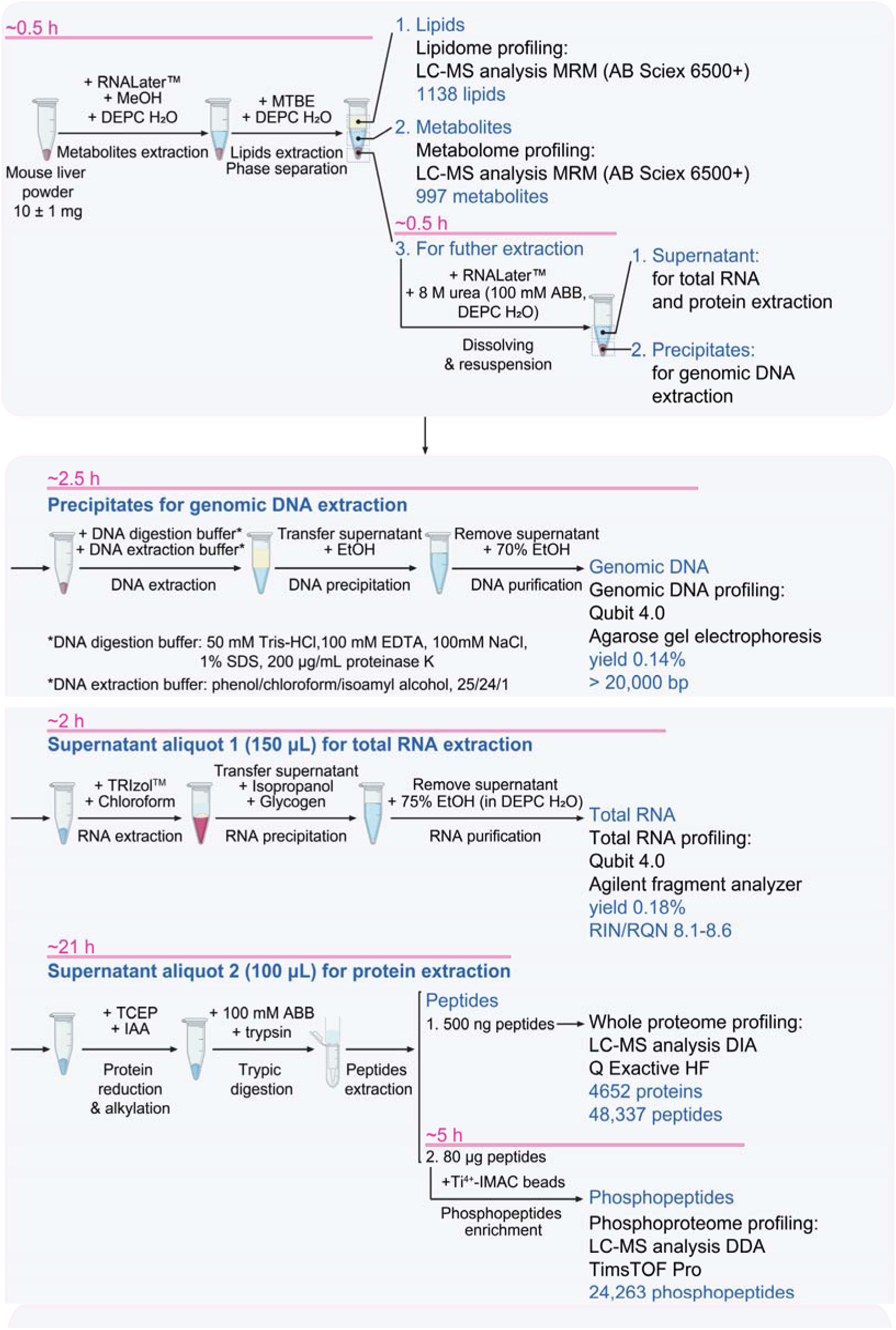
Schematic overview of the Integral-Omics workflow. Metabolites and lipids were initially extracted using the Matyash method, resulting in the identification of 997 metabolites and 1138 lipids. The remaining precipitate was re-lysed in 8 M urea, enabling the extraction of genomic DNA with a 0.14% yield. Further processing of the supernatant led to the extraction of total RNA with a 0.18% yield, and identification of 4652 proteins, 48,337 peptides, 24,263 phosphopeptides, and 18,084 unique phosphorylation sites.

Independent extraction methods and multi-omics extraction kits were used as control groups, and the data showed that sequential extraction did not significantly impact the data for all the omics components. The Integral-Omics workflow yielded comparable omics data to those from independent extractions. Detailed optimization and evaluation data are provided below.

### Evaluation of the metabolome and lipidome analysis

In the first step of the Integral-Omics workflow, we employed two established biphasic extraction methods, namely Folch ^15, 19^ (MeOH/CHCl_3_/DEPC H_2_O, or the Folch method) and Matyash ^16, 17^ (MTBE/MeOH/DEPC H_2_O, or the Matyash method), and a widely used monophasic extraction method ^29, 30^ (MeOH/ACN/DEPC H_2_O, or the MonoP method). We assessed the efficacy of these methods based on the number of identified substances, intra-group reproducibility, inter-group correlation, and its performance to be integrated with subsequent omics extraction. Each method was applied to six biological replicates of mouse liver samples.

These three methods identified a similar number of metabolites (993-999) (**Figure 2A**, **Supplementary Data 1**). However, the MonoP method extracted fewer lipids (1023 ± 14.2) compared to the Folch (1139 ± 0) and Matyash methods (1138 ± 0.4) (**Figure 2B, Supplementary Data 1**). Previous studies suggest lipid and metabolite detection depends on sample amount, extraction methods, and LC-MS settings, with typical identifications ranging from 675 to 1500, consistent with our findings ^31-33^. Venn diagrams showed high overlap in detected metabolites (98.1%) and lipids (91.0%) across methods (**Figures 2C, D**). Pearson correlation analysis confirmed high similarity in metabolite and lipid quantification for Folch and Matyash methods (0.98, 0.99), with MonoP showing lower correlation **(Figures 2E, F)**. Quantitatively, the Matyash method showed the lowest intra-group variance, with median variances for lipids and metabolites below 15%, making it the most reproducibly quantifiable method (**Figures 2G, H**). Considering both qualitative and quantitative data, the Matyash method achieves a balance between identification count and reproducibility.

**Figure 2.**
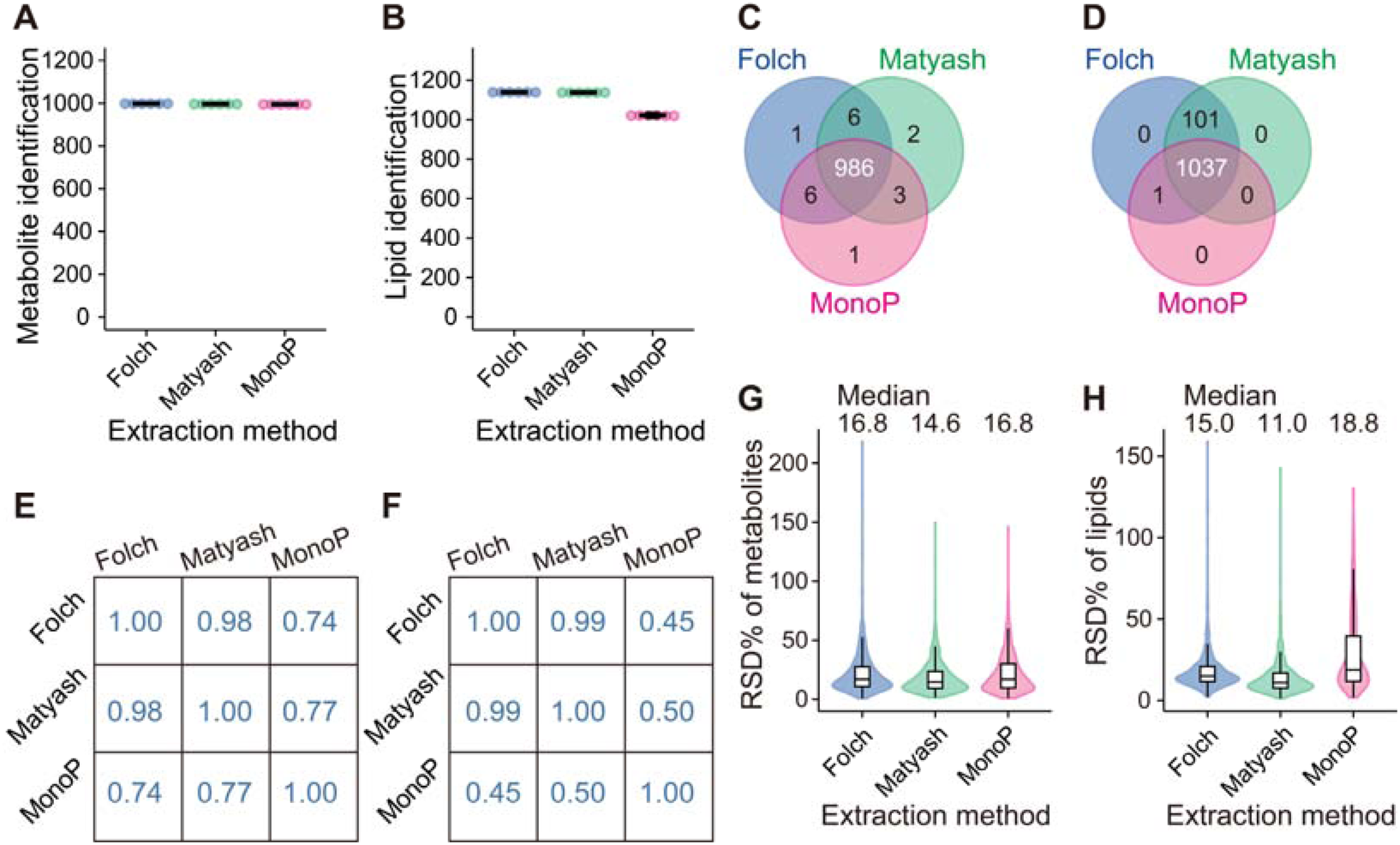
Evaluation of the metabolites and lipids co-extraction part in the Integral-Omics workflow (n = 6 biological replicates per group). **(A)** Number of metabolites identified using the Folch, Matyash, and MonoP methods. **(B)** Number of lipids identified using the three methods. **(C)** Venn diagram showing metabolite overlap among the three extraction methods. **(D)** Venn diagram showing lipid overlap among the three extraction methods. **(E)** Pearson correlation coefficients indicating metabolite similarity across the three methods. **(F)** Pearson correlation coefficients indicating lipid similarity across the three methods. **(G)** Relative standard deviation (RSD%) of detected metabolites across the three methods. **(H)** RSD% of detected lipids across the three methods.

Further analysis of the lipid and metabolite categories obtained by the Matyash method revealed that the metabolome encompassed 16 major metabolite categories and 47 subcategories, while the lipidome included six major lipid classes and 42 subcategories (**Figures S1A, B**). In summary, the Matyash method is the preferred choice among the three co-extraction methods for lipids and metabolites.

### Evaluation of the genome and transcriptome analysis

Following lipidomic and metabolomic extraction using the Folch, Matyash, and MonoP methods, the resulting precipitates were redissolved in 8 M urea (100 mM ABB, DEPC H_2_O) for subsequent omics profiling. After ice bath sonication (4LJ, 40 Khz, 10 min), 100 μL of supernatant was used for protein extraction, 150 μL for RNA extraction via TRIzol™, and the precipitate for DNA extraction via the phenol-chloroform-isoamyl alcohol method ^34, 35^.

The phenol-chloroform-isoamyl alcohol method was used for DNA extraction to assess the reliability of the Integral-Omics workflow, with no significant difference in DNA yield compared to independent extraction (**Figure 3A**). Agarose gel electrophoresis confirmed that DNA extracted using the Matyash method exceeded 20,000 bp, meeting gene sequencing requirements (**Figure S2B**). The results indicated that the upstream omics extraction of lipids and metabolites didn’t affect the yield and integrity of genomic DNA.

**Figure 3.**
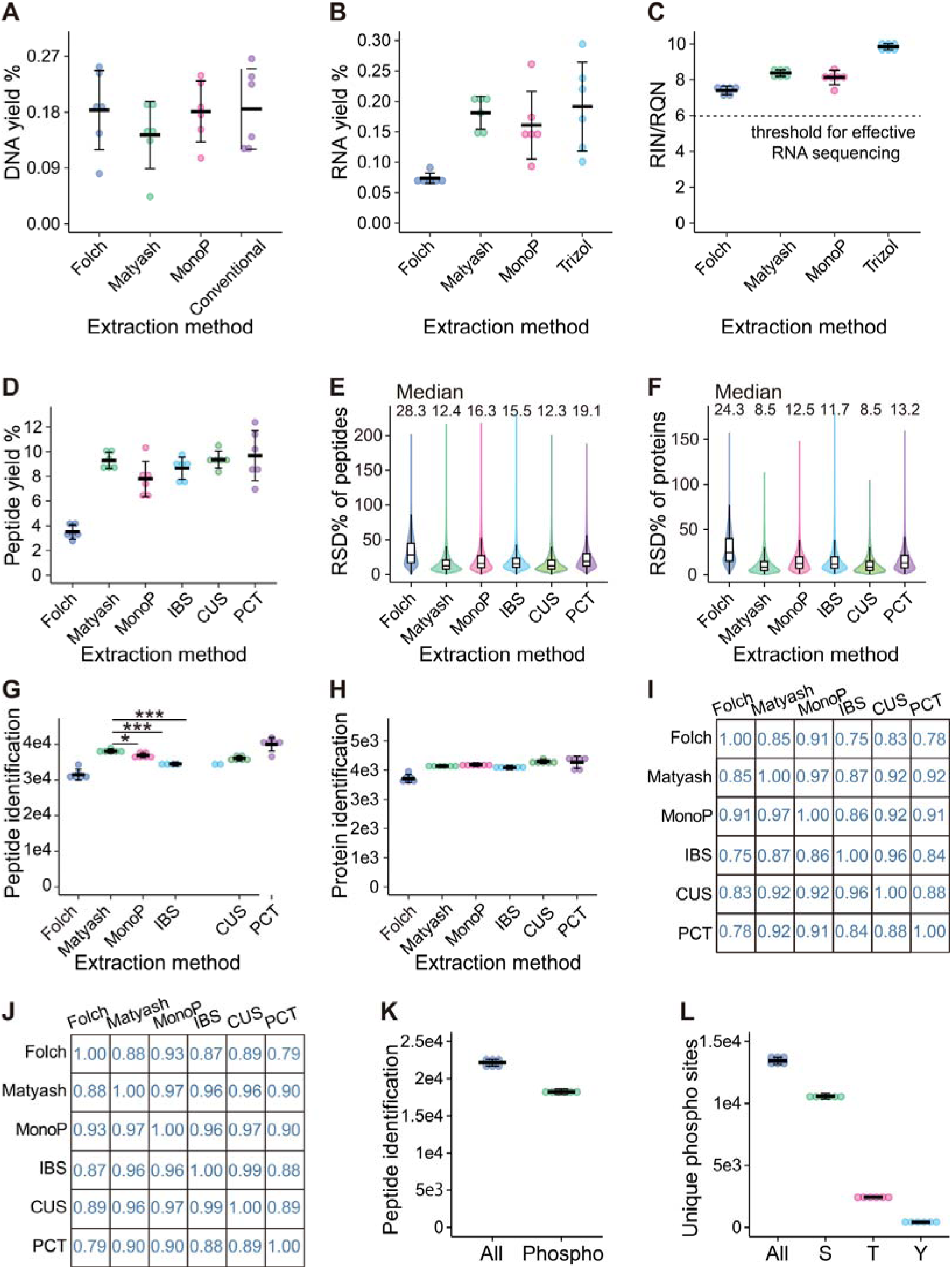
Evaluation of the DNA, RNA, protein and phosphopeptide extraction parts in the Integral-Omics workflow (n = 6). **(A)** DNA yields from the Integral-Omics methods (Folch, Matyash, MonoP) versus independent extraction (Conventional). **(B)** RNA yields from the Integral-Omics methods versus independent extraction (Trizol). **(C)** RIN/RQN values from the Integral-Omics methods compared to the Trizol method. **(D)** Peptide yields from the Integral-Omics methods versus independent protein extraction methods (IBS, CUS, PCT). **(E)** RSD% of detected peptides across the Integral-Omics methods versus independent methods. **(F)** RSD% of detected proteins across the Integral-Omics methods versus independent methods. **(G)** Number of peptides identified in the Integral-Omics methods compared to independent methods (***, p<0.001; *, p<0.05). **(H)** Number of proteins identified in the Integral-Omics methods compared to independent methods. **(I)** Pearson correlation for peptide identification between the Integral-Omics methods and independent methods. **(J)** Pearson correlation for protein identification between the Integral-Omics methods and independent methods. **(K)** Peptides and phosphopeptides identified through the Integral-Omics workflow (Matyash method). **(L)** Number of unique phosphorylation sites identified through the Integral-Omics workflow (Matyash method).

To assess the reliability of transcriptomic data from the Integral-Omics workflow, we compared it with the independent Trizol method. The Folch method yielded significantly lower total RNA and RNA quality (RIN/RQN values) compared to the Matyash and MonoP methods (**Figures 3B, C**). There was no significant difference between the Matyash and MonoP methods in terms of total RNA yields and RIN/RQN values. We further compared RNA yields from the Matyash method to those from the Trizol method, finding them comparable, indicating no detectable RNA loss from upstream omics extraction. All six biological replicates from the Matyash method had RIN/RQN values exceeding 8.1 (**Figure 3C**), meeting transcriptome sequencing requirements (RIN/RQN > 6.0, **Figure S2A**).

In summary, the Matyash-based Integral-Omics workflow does not diminish RNA yield. The upstream omics extraction can induce certain degree of RNA degradation which is acceptable for transcriptome sequencing.

### Evaluation of the proteome and phosphoproteome analysis

Next, 100 μL of the supernatant (tissue lysate in 8 M urea) was used for proteomic extraction. To evaluate the impact of prior metabolomic and lipidomic extractions on the proteome and assess ice bath sonication as a lysis method, we conducted independent proteomic extractions using contact ultrasonication (‘CUS’), Pressure Cycling Technology (‘PCT’), and ice bath sonication (‘IBS’) from the same batch of mouse liver. Each method was tested with six biological replicates. Quantitative protein and peptide data for each group are detailed in **Supplementary Data 1**.

In the Integral-Omics workflow, the Folch method resulted in the lowest peptide yield and identification compared to the Matyash and MonoP methods (**Figures 3D, G, H**). This likely occurred because the sample precipitate in the Folch method remained in the middle layer, leading to sample loss during layer aspiration and negatively impacting protein extraction. Consequently, the Folch method is deemed unsuitable for this workflow. While no significant differences were observed between the Matyash and MonoP methods regarding peptide yield and protein identification, the Matyash method showed higher peptide identification and lower within-group variation, making it the preferred method (**Figures 3E-H**).

Further comparison of the Matyash method with three independent protein extraction methods showed that Matyash’s peptide yield (9.3 ± 0.7%) and protein identification (4138 ± 49) were comparable to the controls (IBS, CUS, PCT), while the peptide count (38,046 ± 636) was significantly higher than in IBS and CUS (**Figure 3G**). Moreover, Matyash demonstrated lower median within-group variation of peptides and proteins than IBS and PCT, comparable to CUS (**Figures 3E, F**). Pearson correlation coefficients indicated a strong correlation (>0.90) between the Matyash method and CUS/PCT in peptide and protein detection (**Figures 3I, J**). Overall, the Matyash method within the Integral-Omics workflow provides similar depth and quantitative accuracy to independent extractions.

Following comprehensive multi-omics analysis, the Matyash method was selected as the optimal approach for the Integral-Omics workflow due to its maintained coverage depth and quantitative accuracy across metabolomics, lipidomics, genomics, transcriptomics, and proteomics. Given these advantages, we employed the Matyash method for downstream phosphoproteomics analysis.

The Matyash workflow identified 28,719 total peptides, with 24,263 being phosphopeptides (**Figure 3K, Supplementary Data 1**), demonstrating highly efficient recovery (84.5%) and identification of phosphopeptides consistent with prior studies ^36^. Additionally, 18,084 unique phosphorylation sites were identified, comparable to findings in Paula *et al.*’s study ^37^ (**Figure 3L)**. The distribution of serine (76.4%), threonine (20.0%), and tyrosine (3.6%) phosphorylation sites aligns with existing literature ^37, 38^.

Thus, the Matyash method offers similar depth and accuracy in proteomic profiling as independent methods and is optimal for the Integral-Omics workflow, including downstream phosphoproteomics analysis.

### Comparison of the Integral-Omics workflow with a multi-omics extraction kit

Currently, no workflow offers sequential profiling of six omics like Integral-Omics. For comparison, we selected the widely used Qiagen AllPrep DNA/RNA/Protein kit, referred to as the ‘AllPrep’ method. Since this kit only extracts DNA, RNA, and proteins, we established the ‘LM_AllPrep’ method, which adds metabolite and lipid extraction using the Matyash method before applying the AllPrep kit. Both methods were tested on the same batch of homogenized mouse liver powder, with six biological replicates per group.

Our results showed that the genomic DNA yield from the Integral-Omics workflow was significantly higher than that from both the AllPrep and LM_AllPrep methods (**Figure S3A**). The Integral-Omics workflow also extracted significantly higher total RNA yield compared to the LM_AllPrep method (**Figure S3B**). Although the RNA yield from the Integral-Omics workflow was higher than that from the AllPrep method, the difference was not statistically significant (p > 0.05) (**Figure S3B**). However, the RIN/RQN values of RNA in the Integral-Omics workflow were slightly lower, probably due to the relatively sophisticated procedures involved (**Figure S3C**). Nevertheless, the RIN/RQN values of the thus extracted RNA samples ranged from 8.6 to 8.9, which are much higher than the threshold for effective RNA sequencing (RIN/RQN of 6.0) (**Figure S3C**).

Our data showed that the Integral-Omics workflow offers significant benefits to proteomic profiling, evidenced by the substantially higher peptide yield and the greater number of identified proteins and peptides in the Integral-Omics workflow compared to both the AllPrep and LM_AllPrep methods (**Figures S3D, E, F**). Furthermore, the peptide yield and the number of identified proteins in the LM_AllPrep method were significantly lower than that in the AllPrep group, indicating that lipid and metabolite extraction by Matyash method adversely impacted the performance of Qiagen AllPrep kit in terms of protein extraction.

In summary, the Integral-Omics workflow outperformed the Qiagen AllPrep kit at the genomic and proteomic levels, although the later RNA extraction step in the Integral-Omics workflow led to slightly higher RNA degradation, still within acceptable sequencing quality.

### Application of Integral-Omics workflow to clinical biopsy tissue specimens

In order to evaluate the feasibility of the Integral-Omics workflow for clinical biopsy specimens, we employed tumor tissues and matched normal adjacent tissues (NATs) from six colorectal cancer (CRC) patients, each with a weight of 10±1 mg. Through this workflow, we extracted metabolites, lipids, genomic DNA, total RNA, proteins, and phosphopeptides. In total, we identified 1116 metabolites, 1920 lipids, 9257 proteins, 132,537 peptides, and 40,107 unique phosphorylation sites (**Supplementary Data 2**). The extracted RNA showed RIN/RQN values ranging from

4.0 to 7.6, which are slightly lower than the mouse liver samples, possibly due to the sophisticated sampling and storage conditions of clinical specimens, as we have previously investigated ^39^. Nevertheless, 92,900 transcripts corresponding to 18,069 genes were detected (**Supplementary Data 2**). Whole exome sequencing (WES) revealed 99.7% exon coverage using GRCh38, with 99.1% of targeted exons covered at over 20-fold depth and an average depth of 207.8X. We identified 253,584 unique SNPs across 30,571 gene symbols and 92,147 unique indels across 19,848 gene symbols (**Supplementary Data 3 and 4**).

Comparing multi-omics data between tumor and NAT tissues, we identified 8579 unique somatic SNVs and 152 somatic indels (**Supplementary Data 5**). The most frequently mutated genes included *APC*, *TP53*, and *KRAS*, all known CRC markers, with *APC* and *TP53* mutations in all patients, consistent with findings from previous CRC cohort studies reported by the TCGA ^40^, Vasaikar *et al.* ^41^, and Kalmar *et al* ^42^ (**Supplementary Data 5**). Additionally, significant upregulation was observed in 55 metabolites, 233 lipids, 266 proteins, 304 phosphopeptides, 1827 genes, and 2112 transcripts in tumors, while 16 metabolites, 170 lipids, 343 proteins, 211 phosphopeptides, 1770 genes, and 1994 transcripts were downregulated (**Supplementary Data 6**). Ingenuity pathway analysis (IPA) identified 95 enriched pathways across all omics levels, including those associated with CRC, such as DNA methylation and transcriptional repression signaling ^43^, autophagy ^44^, AMPK Signaling ^45^, and ferroptosis signaling pathway ^46^ (**Supplementary Data 7**).

Ferroptosis has emerged as a promising target for colorectal cancer treatment ^47, 48^. Our IPA enrichment analysis revealed that ferroptosis is suppressed in CRC tumors. By integrating molecules from both the IPA and KEGG Mapper ferroptosis pathways, we identified one metabolite (L-glutamic acid, upregulated in tumor) and two lipids (PE (18:0_22:4), upregulated in tumor, and FFA (20:4), with no significant difference) related to ferroptosis (**Supplementary Data 8**). Additionally, we identified a total of 66 ferroptosis-related genes, corresponding to 43 proteins, 219 phosphopeptides, and 334 transcripts (**Supplementary Data 8**). Upon analyzing the whole exome sequencing data of these 66 genes, we found 27 mutations exclusively detected in tumor samples (**Supplementary Data 9**). Notably, the TA to T mutation at position 50996580 in the *SLC11A2* gene on chromosome 12 exhibited the highest frequency in tumor samples, reaching 50%. Subsequently, we extracted 29 genes from the KEGG Mapper ferroptosis pathway identified in our dataset and mapped the ferroptosis pathway, as shown in **Figure 4**. Using radar charts, we showed the tumor-to-NAT fold changes (FC) of these molecules at the transcriptome (both gene and transcript levels), proteome, and phosphoproteome levels. Our analysis of ferroptosis pathways revealed key genes that may inhibit ferroptosis and promote tumor survival, including downregulation of *TF* and *FTH1*, potentially reducing iron levels, and upregulation of *SLC3A2* and *SLC7A11*, enhancing GSH synthesis and oxidative stress defense. This suggests CRC tumors evade ferroptosis through multiple mechanisms, contributing to progression.

**Figure 4.**
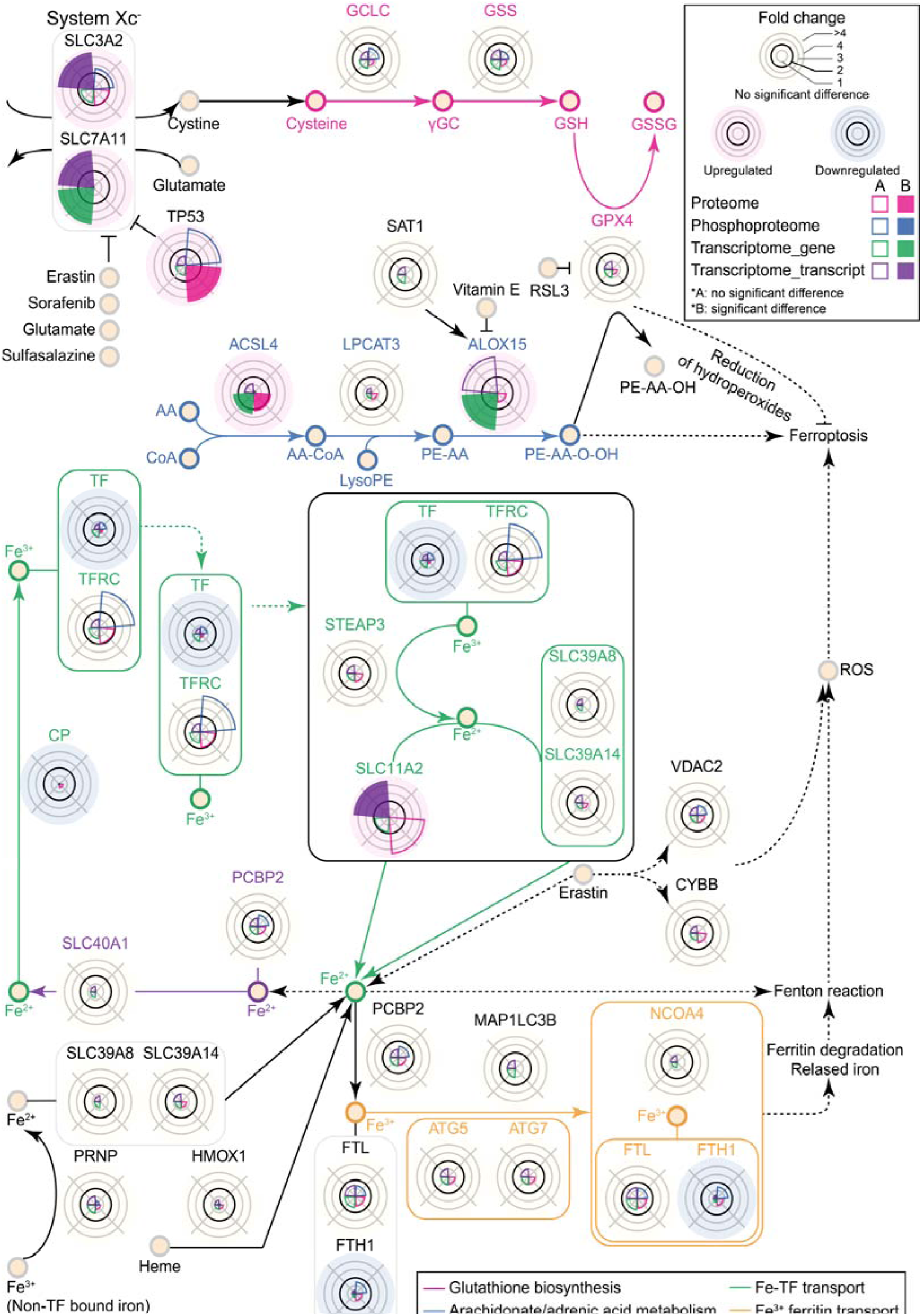
Integrated analysis of ferroptosis pathway in CRC tumors and NATs. This figure presents the ferroptosis pathway, highlighting 29 molecules identified in our dataset. Radar charts show the fold change (FC) of tumors relative to NATs across four omics layers: proteome (pink), phosphoproteome (blue), transcriptome gene level (green), and transcriptome transcript level (purple). Yellow backgrounds indicate no significant difference, pink indicates upregulation, and blue indicates downregulation in tumors. The pathway includes five sub-pathways: glutathione biosynthesis (pink), arachidonate/adrenic acid metabolism (blue), Fe^2+^ ferroportin transport (purple), Fe-TF transport (green), and Fe^3+^ ferritin transport (yellow).

This study demonstrates the Integral-Omics workflow’s ability to extract comprehensive molecular data from biopsy-level tissue specimens, offering insights into regulatory pathways such as ferroptosis.

## Discussion

The Integral-Omics workflow offers a robust and sequential extraction method for metabolites, lipids, genomic DNA, total RNA, proteins, and phosphopeptides from minimal tissue samples. This workflow has been benchmarked against independent extraction methods and existing multi-omics kits, demonstrating that it preserves the integrity and quality of all omics components. Our results confirm that the sequential extraction process does not compromise data quality, producing reliable and reproducible multi-omics data comparable to those obtained through established methods.

A key feature of the workflow is the modified Matyash method for the co-extraction of lipids and metabolites. This method was selected based on a thorough comparison with other extraction methods (Folch and MonoP) due to its ability to minimize degradation risks and its compatibility with subsequent omics extractions. The biphasic extraction effectively separates lipids, metabolites, and nucleic acids, ensuring specific enrichment of each component, while minimizing procedural errors and preserving sample integrity.

The workflow also utilizes 8 M urea in ice bath sonication for the extraction of DNA, RNA, and proteins, a process that maintains nucleic acid integrity and enhances protein identification. To further protect RNA and DNA from degradation, the workflow employs DEPC water, low-temperature processing, and the addition of RNAlater during initial extraction steps. This setup allows for flexibility to selectively extract specific omics components depending on the research needs, making it adaptable for various experimental designs.

Validation using clinical samples from colorectal cancer patients underscores the workflow’s applicability in a clinical research setting, particularly for early-stage disease research where sample sizes are often limited. Despite some RNA degradation and the length of the process (approximately 27 hours), the workflow delivers comprehensive multi-omics data from small biopsy-level samples, making it particularly valuable for studying early-stage tumors and other conditions where sample availability is a limiting factor.

In summary, the Integral-Omics workflow is a powerful tool for multi-omics analyses, capable of providing comprehensive molecular profiling from minimal tissue samples. Its integration of multiple omics extractions within a single workflow not only enhances efficiency but also ensures data quality and integrity, offering significant potential for advancing disease research and personalized medicine strategies. Future work will focus on optimizing the workflow to further reduce processing time, explore its applicability to even smaller tissue samples, and extend its use to a broader range of biological and clinical contexts.

### Materials and Methods Mouse liver

All animal maintenance and experimental procedures were carried out in compliance with Westlake University’s Animal Care Guidelines, and all animal studies received approval from the Institutional Animal Care and Use Committee (IACUC) of Westlake University, Hangzhou, China, under animal protocol #23-072-GTN. Detailed information regarding the extraction of mouse liver is provided in the **Supplementary Note 1**. Mouse liver samples were stored in liquid nitrogen or dry ice, while for long-term storage, samples were frozen at -80°C to preserve biochemical integrity.

### Clinical tissue specimens

Tissue samples were collected from six colorectal cancer (CRC) patients (stages I-III) who underwent surgery in 2023 at Ruijin Hospital, Shanghai Jiao Tong University. Tumor and matched normal adjacent tissues (NATs) were collected 5 cm from the tumor margin. Ethical approval was obtained from the Ruijin Hospital (ID: 2020-384) and Westlake University (20240513GTN002), with informed consent from all patients. Patient details are summarized in **Supplementary Table 1**.

### Metabolite and lipid extraction using Matyash method

To mitigate RNA degradation during metabolome and lipidome extraction, we controlled temperature on ice, used DEPC-treated water, and added 40 µL of RNAlater^TM^ per sample. After adding 225 µL methanol and 35 µL DEPC-treated water, samples were vortexed, followed by the addition of 750 µL MTBE, and incubated on a shaker at 4°C and 1500 rpm for 10 minutes. After adding 187.5 µL DEPC-treated water, samples were phase-separated on ice and centrifuged. The upper layer was for lipidomics, the lower layer for metabolomics, and the bottom precipitate for subsequent DNA, RNA, and protein extraction. Detailed procedures for Folch and MonoP methods are in **Supplementary Note 2**.

### Genomics DNA, total RNA and protein extraction, and phosphopeptides enrichment

After metabolite and lipid extraction, the precipitate was used for further omics extractions. We added 40 µL RNAlater^TM^ and 500 µL 8 M urea lysis solution (100 mM ammonium bicarbonate in DEPC-treated water), followed by ice bath sonication at 40 kHz for 10 minutes. After sonication, the samples were placed on a shaker and incubated at 4°C and 1500 rpm for 15 minutes. Following incubation, a 100 μL aliquot of the sample was utilized for subsequent protein extraction. Then sample was subjected to centrifugation in a high-speed refrigerated centrifuge at 4°C and 12,000 rpm for 10 minutes. A 150 μL aliquot of the supernatant was utilized for subsequent total RNA extraction, and the remaining sample precipitate was preserved for genomic DNA extraction. DNA was extracted using the phenol-chloroform-isoamyl alcohol method ^35^, RNA by the Trizol method ^34^, and proteins were processed for proteomics and phosphopeptide enrichment as described by Zhou et al ^49^.

### Integral-Omics workflow

Refer to **Supplementary Note 3** for a detailed description of the Integral-Omics workflow. Materials and reagents are listed in **Supplementary Table 2**.

### Independent Extractions

Genomic DNA, total RNA, and proteins were independently extracted from the same batch of mouse liver powder (10 mg per sample, n = 6) using standard methods as control groups: phenol-chloroform-isoamyl alcohol for DNA (referred to as the Conventional group), Trizol for RNA (referred to as the Trizol group), and three proteomic extraction methods (referred to as the IBS group, CUS group, PCT group) described in **Supplementary Note 4**.

### Multi-omics extraction by Qiagen AllPrep DNA/RNA/Protein Mini kit

For the AllPrep group, DNA, RNA, and proteins were extracted according to the Qiagen AllPrep DNA/RNA/Protein kit (Qiagen, Hilden, Germany, 80004) protocol. In the LM_AllPrep group, metabolites and lipids were first extracted using the Matyash method before applying the Qiagen kit for DNA, RNA, and protein extraction.

### Lipidomics and metabolomics data acquisition, qualitative and quantitative analysis

Samples were thawed on ice, and internal standards were added before LC-MS analysis. Lipidomic analysis involved a 20-minute gradient and specific analytical conditions, while metabolomic analysis used a 14-minute gradient. Data were analyzed using MWDB, Multi Quant software, and visualized in Rstudio (**Supplementary Note 5**).

### Proteomics and Phosphoproteomics Data Acquisition, Identification, and Quantification

For both proteomics and phosphoproteomics analyses, peptides (500 ng for mouse liver, 250 ng for colorectal cancer) and phosphopeptides were separated using 75 µm × 150 mm C18 analytical columns in HPLC nanoflow systems with gradient elution. Quantification was performed using Orbitrap (Thermo Q Exactive™ HF) mass spectrometers in DIA mode, and timsTOF Pro (Bruker Daltonics, Germany) mass spectrometers in DIA-PASEF and DDA-PASEF modes. Detailed methodologies are provided in the **Supplementary Note 6**.

### mRNA library preparation, sequencing, and data analysis

mRNA was enriched, fragmented, reverse transcribed to cDNA, and prepared for sequencing on the DNBSEQ platform. Clean reads were aligned to gene sets using Bowtie2, with expression analysis by RSEM and differential expression by DESeq2 (Supplementary Note 7).

### Whole exome sequencing library preparation, sequencing, and data analysis

Genomic DNA was fragmented, end-repaired, and ligated with adapters for sequencing. Clean reads were mapped to the human genome using BWA, with variant calling and annotation performed by GATK (**Supplementary Note 8**).

## Data availability

Mass spectrometry proteomic and phosphoproteomic raw data are deposited in the ProteomeXchange Consortium via iProX under accession PXD054560. Transcriptome and whole exome sequencing raw data are available in the Genome Sequence Archive (GSA) under accession HRA007600 and PRJCA026610. Supplementary Information and Source Data file contain essential raw and processed data.

## Acknowledgments

This work is supported by grants from the National Key R&D Program of China (Grants No. 2022YFF0608403, 2021YFA1301600, 2020YFE0202200 and 2019YFA0801800), the “Pioneer” and “Leading Goose” R&D Program of Zhejiang (2024SSYS0035), and the Westlake Educational Foundation, Shanghai Science and Technology Commission (20JC1410100). The language of this manuscript has been polished by ChatGPT (v 4o) with human oversight and control. Figure 1 was created through Biorender.com. The authors would like to express their gratitude to the patients who participated in this study for their willingness to share clinical samples. We thank Mingrun Xuan, Dr. Cheng Zhang, and Guojin Lin for their valuable insights and assistance.

## Author contributions

W.L. and T.G. conceived and designed the project. W.L. conducted the experiments, analyzed data, and wrote the manuscript. J. Z., J. S. and Y. W., procured the clinical samples, collected the patient information and examined the pathology of the samples. R.S. assisted in manuscript writing. T.G. and Y.Z. supervised the project and revised the manuscript. All the authors read and approved the manuscript.

